# Plis de passage in the Superior Temporal Sulcus: Morphology and local connectivity

**DOI:** 10.1101/2020.05.26.116152

**Authors:** C. Bodin, A. Pron, M. Le Mao, J Régis, P. Belin, O. Coulon

## Abstract

While there is a profusion of functional investigations involving the superior temporal sulcus (STS), our knowledge of the anatomy of this sulcus is still limited by a large variability across individuals. Several “plis de passage” (PPs), annectant gyri buried inside the fold, can separate the STS into distinct segments and could explain part of the observed variability. However, an accurate characterization is lacking to properly extract and fully understand the nature of PPs. The aim of the present study is twofold: *i.* to characterize the STS PPs by directly identifying them within individual STS, using the geometry of the surrounding surface and considering both deep and superficial PPs. *ii.* to test the hypothesis that PPs constitute local increases of the short-range structural connectivity. Performed on 90 subjects from the Human Connectome Project database, our study revealed that PPs constitute surface landmarks that can be identified from the geometry of the STS walls and that they constitute critical pathways of the U-shaped white-matter connecting the two banks of the STS. Specifically, a larger amount of fibers was extracted at the location of PPs compared to other locations in the STS. This quantity was also larger for superficial PPs than for deep buried ones. These findings raise new hypotheses regarding the relation between the cortical surface geometry and structural connectivity, as well as the possible role of PPs in the functional organization of the STS.

## Introduction

Neuroanatomists of the 19th century were closely interested in the origin but also the individual variability of cortical folding. They noticed in this context that folds could be subdivided into sub-units separated by specific morphological landmarks. The term “pli de passage” (PPs) was first introduced by the anatomist Gratiolet (1854) in reference to inter-connecting gyri buried inside the main furrows and causing a protrusion at the bottom. In his book comparing the cerebral sulci of primates, Gratiolet points out that PPs can share similar patterns of organization across species and therefore constitute markers of interest. Later on, Broca (1888) re-used this term and reported the existence of three transverse gyri along the central sulcus (CS) connecting the pre- and post-central gyri: the PPs frontal superior, middle and inferior. Such subdivision of sulci into several pieces is made clearer by Cunningham’s pioneering work on cortical development: he described how folds appear first as distinct segments that merge at a later stage in parallel with cortical expansion (Cunningham 1890a; 1890b; 1897). The central sulcus, in particular, would originate from two pieces separated by what he called a “deep annectant gyrus” (Cunningham 1897) characterized either by a clear elevation of the fundus or a thickening of its two interlocking extremities which can unite at the bottom (Cunningham 1892; White et al. 1997). This landmark, that would correspond to Broca’s “PPs fronto-parietal moyen” (PPFM), can persist until adulthood leading to a segmented aspect of the sulcus (Cunningham 1897; Regis et al. 2005; Cykowski et al. 2008). More recently, it was shown to reflect the position of the hand motor area (Boling et al. 1999; Cykowski et al. 2008; J.-F. Mangin et al. 2019) also co-localized with the omega-shaped bending of the surface or “hand knob” (Yousry et al. 1997).

PPs can constitute landmarks for a better understanding of the inter-individual variability. Two complementary approaches have provided evidence in this regard. While the first approach tends to separate the population into distinct anatomical patterns, the second one tends to identify what are the common features to all individuals. In relation to the former, Ono, Kubik, and Abernathey (1990) used the term “sulcal interruptions” to describe the various folding patterns and their relative proportion across individuals. Based on their depth level, they were classified into “true” interruptions causing a clear discontinuity and “pseudo interruptions” generating a slight deformation of the fundus. This qualitative description of PPs can be found in more recent studies, sometimes under other names such as “submerged gyri” or “gyral/cortical bridges”. Especially, they were shown relevant to describe the anatomo-functional organization across individuals in the intra-parietal (Zlatkina et Petrides 2014), cingulate (Paus et al. 1996; Amiez et al. 2013), collateral (Huntgeburth et Petrides 2012) and post-central (Zlatkina et Petrides 2010; Zlatkina, Amiez, et Petrides 2016) sulci. This first approach provides a direct insight into the variability of cortical folding, but it is limited by manual identification procedures. The second approach is well illustrated by the “sulcal roots” model (Régis et al. 1995; Regis et al. 2005), which advocates a common organization scheme of folds across individuals. Like a map, all sulcal roots can be arranged on a flat representation of the cortical surface (Auzias et al. 2013), each of them being delimited by two parallel and two orthogonal gyri (corresponding to transverse PPs locations). This putative organization may be visible at the fetal stage when folding starts to appear and then changes during gyrification to eventually lead to inter-individual variability as observed after birth, in line with Cunningham’s early ideas. This model also gave rise to the study of the anatomical landmarks dual to the PPs, the ‘‘sulcal pits’’, which are points of maximum depth within folds showing strong inter-subject reproducibility (Im et al. 2010; G. Auzias et al. 2015).

What emerges from this literature however is a lack of a clear definition of PPs. It is not yet clear whether they are inherently present in a stable pattern as suggested by the root-based emergence of sulci (Cunningham 1890a; 1890b; 1897; Regis et al. 2005) or whether different patterns exist in individuals (Ono, Kubik, et Abernathey 1990). One major question that arises here is whether or not PPs should be selected on the basis of the sulcal depth. In other words, can we consider them to vary from being completely apparent at the surface to being completely buried such that they are not related to any decrease of sulcal depth. In order to investigate this, we need to be able to provide a novel anatomical characterization that does not discard the deepest PPs. Such approach, less conservative, was already shown to increase the number of PPs detected while improving the correspondence with motor functions in the post-central sulcus (Zlatkina, Amiez, et Petrides 2016). In this case, the authors were able to identify smaller PPs raising the fundus by only 1 mm and characterized by an unusual curvature of the sulcus.

The case of the STS is interesting regarding the complexity of its anatomo-functional organization. It endorses a rich set of functions, mainly in the perception and processing of social stimuli derived from multiple sensory modalities (Hein et Knight 2008; Lahnkoski et al. 2012; Deen et al. 2015), distributed along its antero-posterior axis (Beauchamp 2015). According to the sulcal roots model, the STS axis would be interrupted by six highly reproducible PPs albeit with a varying depth (Ochiai et al. 2004). This fixed number is less obvious through visual observation, where the number of PPs was shown to vary from 0 to 4 with a greater amount in the left STS (Ono, Kubik, et Abernathey 1990). This asymmetric distribution was shown to interfere with a global asymmetry of the STS depth, referred to as the “STAP” (Superior Temporal Asymmetrical Pit) (Leroy et al. 2015; Le Guen, Leroy, et al. 2018). In these last studies, PPs were identified as local minima on the two-dimensional depth profile of the sulcus and only those below a certain threshold on their absolute depth value were selected as true PPs. This method appears more restrictive and arbitrary than previous descriptions of the STS patterns (Ono, Kubik, et Abernathey 1990) or root-based organization (Ochiai et al. 2004) and potentially lose information by discarding the most buried PPs.

Finally, some indicators in the literature suggest that PPs could be related to the underlying structural connectivity. Notably, Leroy et al. (2015) reported no STS depth asymmetry (STAP) in people with agenesis of the corpus callosum, for which inter-hemispheric connectivity is severely reduced. In addition, the frequency of PPs was reduced in this group and more symmetrically distributed compared to other groups. In the case of the central sulcus (CS), dense “U-shape” fibers were found at the location of the hand-knob (Catani et al. 2012; Magro et al. 2012; Pron et al. 2018) which contains the “pli-de-passage fronto-parietal moyen” (PPFM) (Broca 1888; Boling et al. 1999). Hence, PPs could be associated with specific structural connectivity patterns and particularly with short-range bundles connecting the two banks of sulci (Catani et al. 2012; P. Guevara et al. 2012; Zhang et al. 2014; Román et al. 2017; Pron et al. 2018). By applying clustering methods to whole-brain tractograms generated from diffusion MR data, two studies in particular have described the U-shape connectivity (Zhang et al. 2014; M. Guevara et al. 2017). They revealed several distinct bundles joining the pre- and post-central gyri at several locations along the central sulcus. Concerning the STS, only few bundles were found to connect its two adjacent gyri, mostly restricted to the posterior portion. Improvements could however be expected from techniques such as Diffusion Spectrum Imaging (DSI), as for instance shown in (Zhang et al. 2014). Knowing the extreme variability of the STS region, methods that take into account the local anatomy may be more appropriate than whole-brain approaches. In Pron et al. (2018), for example, the tractograms were filtered to extract only the short fibers joining the two CS adjacent gyri.

In this context, the goal of the present study was twofold: – First, providing a morphological characterization of the STS plis de passage, using the geometry of the surrounding surface, such that it reveals both superficial and very deep PPs. We assumed here that PPs are distributed on a continuum of representations of the cortical relief, from a clear emergent interruption of the sulcus to a completely buried configuration with no depth variation at the fundus. The literature already suggested several morphological clues to identify them, mostly an elevation of the fundus but also an unusual curvature (Zlatkina, Amiez, et Petrides 2016) and a close interlocking of the two surrounding gyri (Cunningham 1892) for deep PPs. – Second, demonstrating that this morphological characterization is associated with a specific short-range structural connectivity of the STS. We make the hypothesis that PPs constitute places of particular U-shape connectivity connecting the two banks of the STS, similar to what has been found for the central sulcus. Because we consider deep and superficial PPs as part of the same continuum, we expect to find this specific U-shape connectivity under both types of PPs.

## Methods

### Subjects

We used a subset of the Human Connectome Project (HCP) S900 release, for which detailed information is available here: https://www.humanconnectome.org/study/hcp-young-adult/document/900-subjects-data-release. We selected subjects having completed the full diffusion and structural acquisitions, being non twins, right-handed and between 22 and 40 years old. From these criteria, 100 subjects (50 females; 50 males) were randomly sampled in order to obtain an identical age distribution between gender groups. Ten subjects presented potential anatomical abnormalities as noticed in the QC_Issue file of the HCP 1200S release and were then replaced by 10 new subjects meeting the same selection criteria. This subset was then split into two parts: 10 subjects were used for training in the manual identification of PPs in a consistent manner from one individual to another. After the training phase, structural and diffusion data of the remaining 90 subjects (44 males, mean age 28.9 yo) were analyzed and are presented in this paper.

### Image acquisition

Data taken from the HCP database were acquired as follows: structural images were acquired using a modified version of Siemens Skyra 3T scanner (Siemens, Erlangen, Germany) with a maximum gradient strength of 100mT/m, slew rate of 200 T/m/s (reduced to 91T/m/s for diffusion due to peripheral nerve stimulation limits) and a 32-channel head coil. T1-weighted images were acquired using 3D MPRAGE sequence (TR/TE = 2400/2.14ms, flip angle = 8, FOV = 224×224 mm^2, resolution = 0.7 mm isotropic).

Diffusion-weighted images were acquired with a spin-echo EPI sequence consisting of 3 shells of 90 diffusion-weighted volumes each (b=1000, 2000 and 3000 s/mm2) and 6 interleaved b0 volumes (TR/TE = 5520/89.5ms, resolution: 1.25 mm isotropic, FOV = 210×180 mm2, 111 axial slices, multiband factor = 3, partial Fourier = 6/8, echo spacing = 0.78ms). Gradients directions were sampled over the entire sphere, using the electrostatic repulsion method. The entire diffusion sequence was repeated twice with RPE (L->R, R->L).

Structural and diffusion data served as input to our main analysis pipeline **(Figure 1)** designed to extract the connectivity associated with the STS PPs and detailed in the next sections.

**Figure 1:**
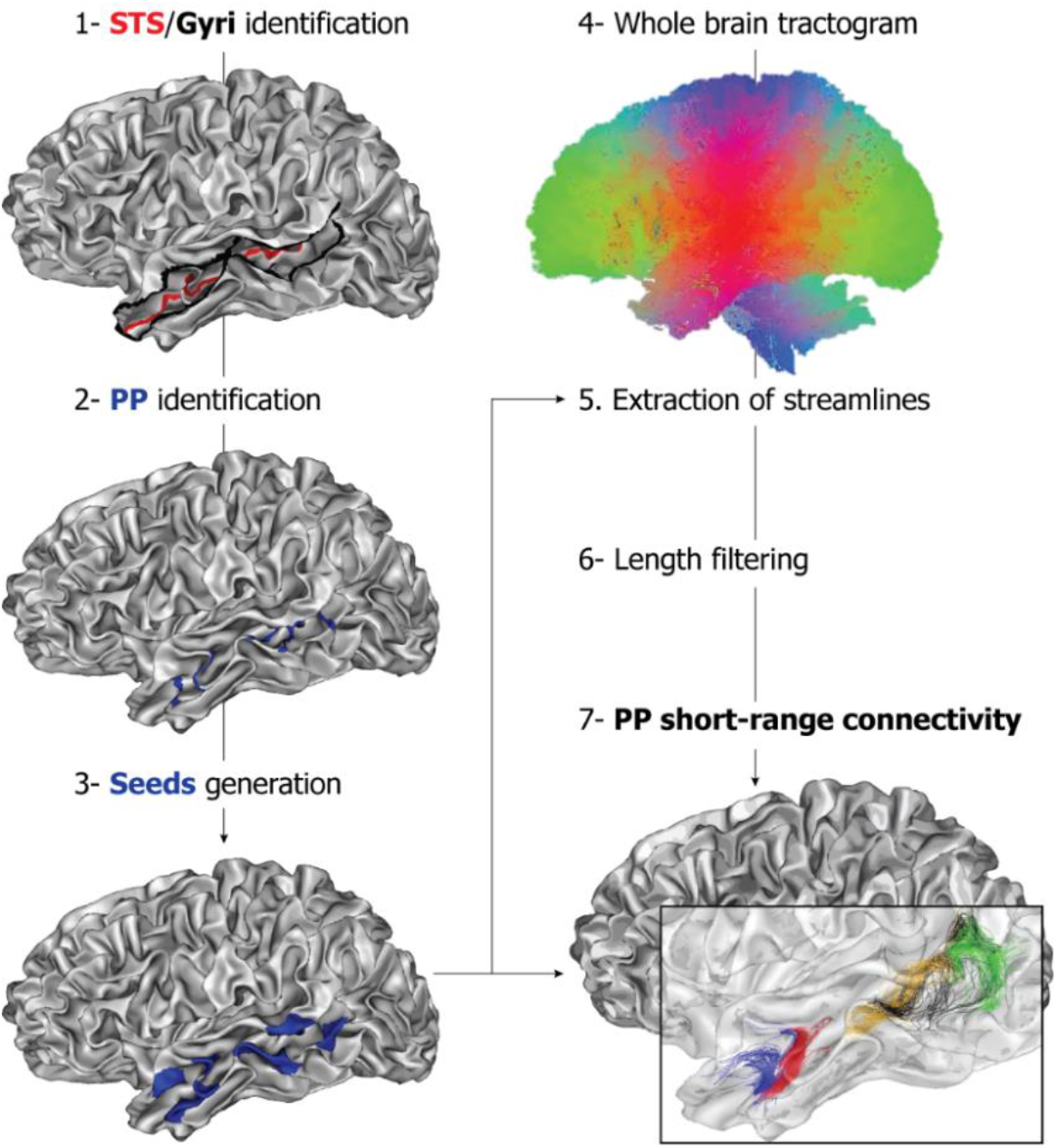
Analysis pipeline for U-shape fibers extraction illustrated for one subject in the left hemisphere. Delimited in length by the STS and in width by the adjacent gyri (1), all PPs are manually identified and drawn (2) on the individual’s surface. Their extremities are used as seeds (3) to extract the underlying short-range connectivity (7). Each colored bundle corresponds to the streamlines extracted from one pair of seeds (one PP).

### Image preprocessing

#### Anatomical images and related maps

Individual T1-images were first segmented using Freesurfer [https://fsl.fmrib.ox.ac.uk/fsl/fslwiki] then imported into the Morphologist pipeline of the BrainVisa (BV) software [http://brainvisa.info] (J. F. Mangin et al. 2004) in order to produce triangular meshes of the grey/white matter interface for both hemispheres of all subjects (example subject **figure 1**). These surfaces will be further referred to as “white mesh”.

Then, we generated depth maps for each individual surface by using the depth potential function (DPF) (Boucher, Whitesides, et Evans 2009), as already done in (Auzias et al., 2015). It is known to provide a regularized estimation of the sulcal depth that takes into account information from both convexity and curvature. Importantly, it was also shown independent of brain size and therefore does not require a normalization procedure (G. Auzias et al. 2015). DPF measure can be either negative or positive depending on whether the vertex is superficial or located in the depth of a sulcus (**top of figure 3**).

Finally, we generated mean curvature maps for each individual white mesh using a finite element method as implemented in BrainVisa. Vertices in the STS fundus appear with minimal curvature while gyri crowns appear with maximal curvature (**top of figure 3**).

#### Diffusion MR images

Diffusion MRI scans preprocessed by the HCP, i.e. corrected for subject movement, susceptibility induced artifacts, eddy-current distortions and diffusion gradients non linearities (Jenkinson et al. 2012; Glasser et al. 2013) were used to build whole brain tractograms with the Mrtrix software (www.mrtrix.com) (J.-D. Tournier, Calamante, et Connelly 2012). Preprocessed scans were first corrected for non-uniform intensity using the ANTS (https://github.com/ANTsX/ANTs) N4 bias correction algorithm (Tustison et al. 2010). For each subject, a multiple shell multiple tissue (MSMT) (cerebrospinal fluid, grey matter, white matter) response function was then derived from the FreeSurfer tissue segmentation using the *dwi2response* command with default parameters. The obtained response was used to fit a constrained MSMT spherical deconvolution model (Jeurissen et al. 2014) on the brain diffusion signal. Whole brain probabilistic tractography (J. D. Tournier, Calamante, et Connelly 2010) was performed using the tckgen command (algorithm = iFOD2, step = 0.625 mm, angle = 45°, nb_streamlines = 5 × 10^6^) with seeding from the GMWM interface volume and anatomical constraints (Smith et al. 2012) stemming from FreeSurfer tissue segmentation. The resulting tractograms were filtered within the Convex Optimization Modeling for Microstructure Informed Tractography (COMMIT) framework (https://github.com/daducci/COMMIT) (Daducci et al. 2015) to remove spurious or overrepresented streamlines and insure the tractogram fitted the diffusion signal. Stick Zeppelin Ball (Panagiotaki et al. 2012) with default diffusivity parameters (parallel diffusivity = 1.7 × 10^−3^ mm^2^.s^−1^, intracellular fraction = 0.7, isotropic diffusities = 1.7 × 10^−3^ and 3.0 × 10^−3^ mm^2^.s^−1^) was used as a forward model. Filtered tractograms **(figure 1.4-**) contained on average one million streamlines. Endpoints of the remaining streamlines were then projected onto the vertices of the GMWM mesh of both hemispheres by minimizing the Euclidean distance.

### Manual identification of landmarks

#### STS and surrounding gyri

The superior temporal sulcus (STS) separates the superior (STG) from the middle temporal gyrus (MTG) in the temporal lobe. The STS fundi and STG/MTG crests of each subject were drawn semi-automatically on their white meshes (**figure 1.1-**)) using the SurfPaint module of the Anatomist visualization software (Arnaud Le Troter, Rivière, et Coulon 2011). We determined manually the anterior and posterior extremities based on anatomical landmarks identifiable in each subject as described in a previous study (Bodin et al. 2017). The anterior extremity was chosen at the tip of the temporal lobe excluding the last polar sulcus that is often oriented transversally to the STS (Ochiai et al. 2004). The posterior extremity was chosen at the intersection between the STS horizontal main branch and its posterior ascending branches (Segal et Petrides 2012). The three lines corresponding to the STS fundus, STG and MTG crests were then drawn automatically, following the deepest (STS) or shallowest (Gyri) path between their respective extremities (A. Le Troter, Auzias, et Coulon 2012).

#### Plis de passage

We identified individually all PPs connecting the two gyri crests (STG, MTG) and crossing the STS fundus line. This identification was first carried out on an independent pool of ten subjects for training. After being able to consistently identify PP locations in a reproducible manner, we drew them on the 90 subjects described in this study. Their extremities were defined as the intersection between the PP apparent crest and the two adjacent gyri lines in SurfPaint. PP lines **(figure 1.2-**) were then automatically drawn between these two extremities as the shortest path maximizing the DPF (A. Le Troter, Auzias, et Coulon 2012). We carried out the identification of PPs according to several morphological criteria that are illustrated in the following **figures 2 and 3**.

**Figure 2:**
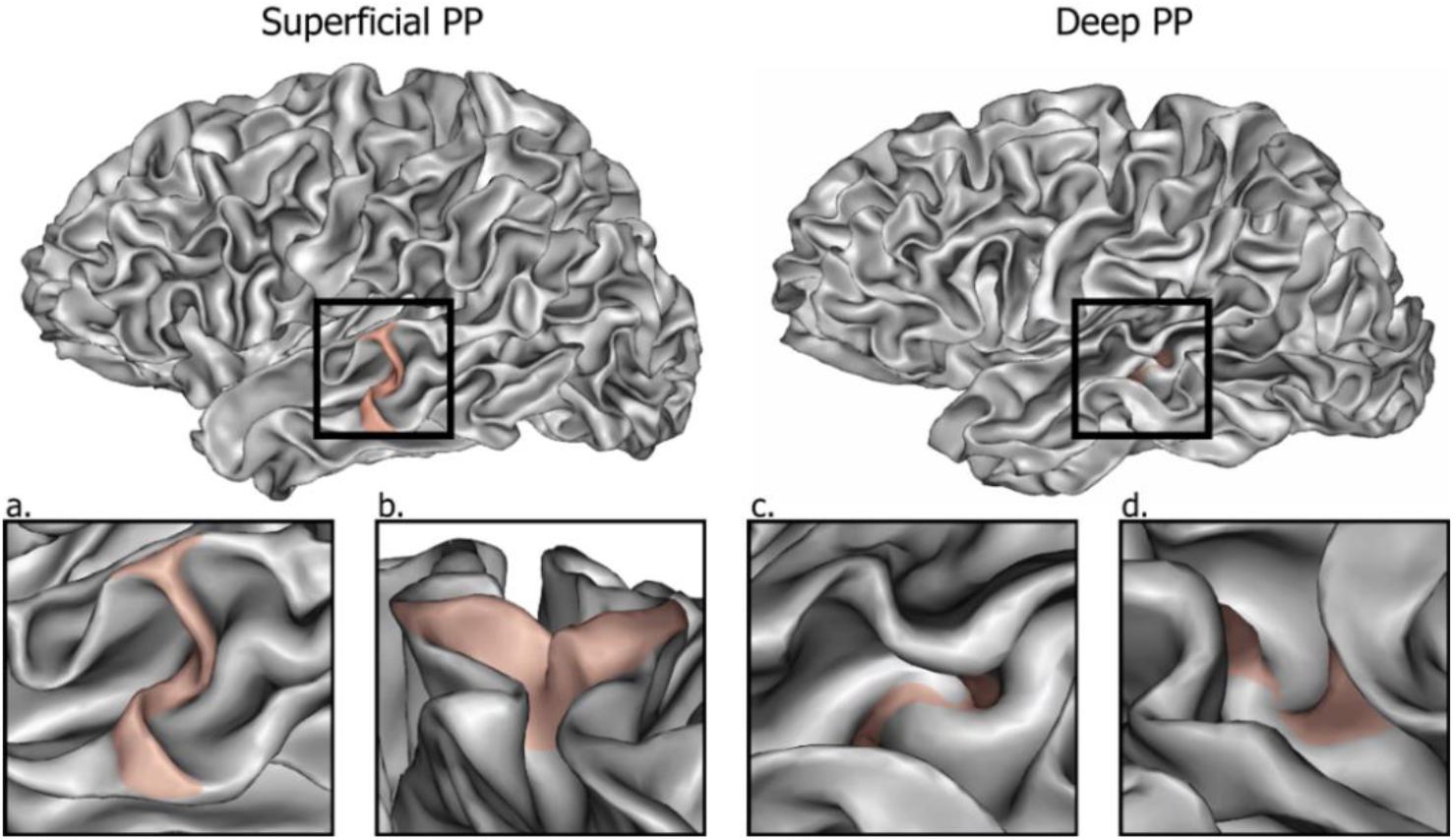
Local cortical morphology observed for superficial and deep PPs (colored areas) illustrated in two example subjects. Only the superficial PP causes a clear separation of the main STS furrow. However, a thorough observation (from top: a,c or laterally: b,d) reveals a pinching of the adjacent walls in both cases, whose visibility is a matter of degree.

**Figure 3:**
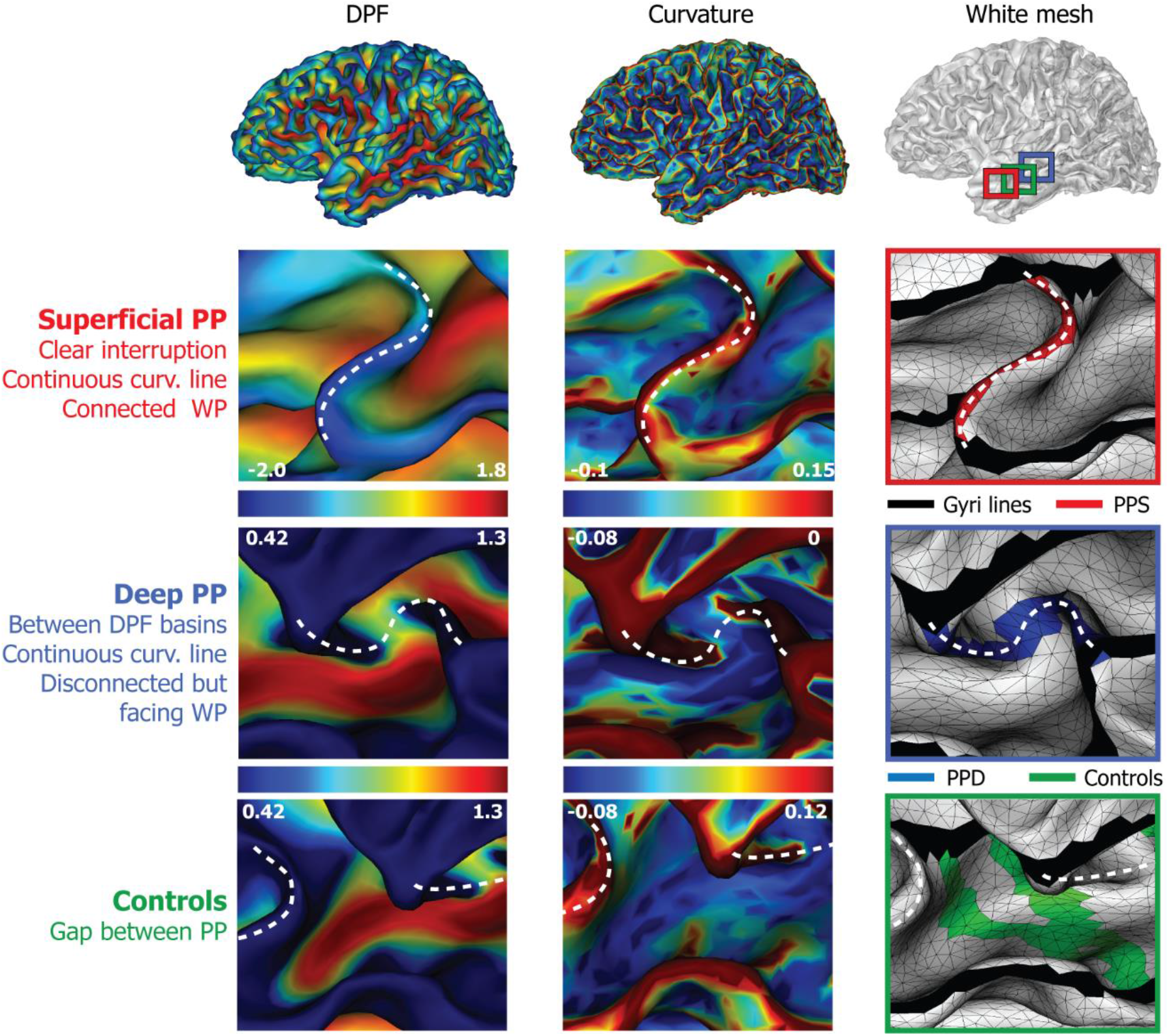
Methods for PPs identification, illustrated for one subject in the left hemisphere. **Top panel:** Surface maps generated in BrainVisa and projected on the white matter surface. DPF map shows an increasing depth from blue to red color. Curvature (finite element method) map shows an increasing curvature from blue to red color. **Squared panels:** Morphological criteria used to characterize PPs (represented by white dotted lines). Zoom windows illustrate the manual drawing of one example superficial (PPS in red) and deep PP (PPD in blue) as well as controls (in green) generated automatically. Once detected based on the morphological criteria (left side), PPs are drawn by selecting the intersections between their two extremities and the adjacent gyri crests (in black). A line is then automatically generated between this pair of vertices passing by the shortest past and constrained by minimal DPF values. Controls are generated using a similar procedure except that their position is generated automatically into the gaps separating PPs locations. If there is enough space, several controls are generated with different orientations. White numbers below color bars indicate the DPF and curvature thresholds employed for each representation. *DPF: Depth Potential Function; PP: plis de passage; WP: wall pinches*.

#### Morphological criteria

As mentioned above, the binary classification of PPs as “present” or “absent”, based on STS depth variations alone (as in Leroy et al. 2015; Le Guen, Leroy, et al. 2018) is insufficient to detect PPs, and to map their topography, namely the spatial arrangement of the different PPs and their respective depth levels (Ono, Kubik, et Abernathey 1990; Ochiai et al. 2004). Indeed, identification based on a two-dimensional depth profile is not appropriate to capture all PPs as it discards the deepest ones. Crucially, PPs are three-dimensional structures whose cortical deformations can be located on the walls of the STS and thus “missed” by the depth profile trajectory. This last point is illustrated in the **figure 2**, showing that all PPs can be characterized by a local deformation, a “pinching” of the STS walls, which is clearly observable for superficial PPs but also true for the deepest ones although to a lesser degree (Cunningham 1890b; White et al. 1997; J.-F. Mangin et al. 2019), and establishes a continuum from a superficial apparent transverse gyrus to a completely buried PP with only wall deformations left.

From these considerations, we further identify all potential PPs characterized by this local deformation and using precise morphological criteria that are described below.

In order to detect PPs, we used DPF, curvature and the white matter mesh previously generated for each hemisphere **(top of figure 3**). These individual maps were carefully examined while varying the DPF and curvature thresholds to highlight distortions of the cortical sheet. Several criteria were used to characterize PPs, as illustrated in one example subject in **figure 3**.

We observed numerous PPs that clearly interrupt the STS transversely, characterized by a strong decrease of the DPF within the sulcus and a high mean curvature forming a thick and continuous line on the surface. We refer to them as “superficial PPs” (in red). A majority of PPs, however, were less visible at first glance and needed more thorough observations based on variable thresholds of the DPF and curvature maps. We refer to them as “deep PPs” (in blue). We selected those that delimitate DPF basins and for which the mean curvature forms a continuous line between the opposite banks of the STS. An additional morphological criterion was the shape of the white mesh at these locations. The presence of a PP was always associated with a pinching of the adjacent sulcal walls, forming a more or less prominent angle on the wall. This can be also observed on the DPF and curvature maps in the form of a superficial crest and a convex angle respectively. Subsequently, a PP was defined as delimited by two opposite “wall pinches” (WPs) facing each other or presenting a slight offset to each other. We hypothesize that WPs of deep PPs are of the same nature than those observed for superficial PPs and that the former constitute a less connected variant of the latter (**figure 2**). If the two WPs are well connected each other they constitute a superficial PPs, whereas if they are not (or too deeply) connected they constitute a deep PPs.

#### Bifurcations

In some cases we observed two PPs sharing one of their wall ‘pinching’. Instead of a single transverse interruption characterized by 2 opposite points, this lead to a triangular “V” (if they join on the MTG) or “Ʌ” (if they join on the STG) shape that we call a “bifurcation”. These structures were not observed in all subjects and often involved one deep and one superficial PP.

#### Controls

In order to test our hypothesis of a short-range connectivity specific to the PPs as we define them, we finally generated “control PPs” as random pairs of points located in the gaps between true PPs (in green **figure 3**). We wanted controls to have a random orientation and location, while being from true PPs. Therefore, when the gap between two consecutive true PPs was at least 15 vertices long, we drew random points on each of the two gyral lines within this gap, with the constraint that each of the two points was at least 5 vertices away from the two PPs. A geodesic shortest path was then computed between the two points and served as a control PP. Up to 3 control PPs were randomly drawn between two true PPs if the above mentioned conditions could be fulfilled. Control PPs were then post-processed exactly like the true PPs.

### U-shape fibers extraction

The main question of the study is whether the morphological definition that we propose for PPs is associated with the location of specific local short-range connectivity, connecting the two banks of the STS. To test this hypothesis, the two extremities of each PP, i.e. intersections between PP lines and gyri crests, were used to define pairs of seeds that were in turn used to extract short-range streamlines (see **figure 1**). The seeds (**3-**) were built as geodesic circles with a given radius around each intersection. A range of radius was used, from 8 to 16 mm. Each individual tractogram was then filtered to extract only the streamlines that connect pairs of PPs seeds (**5-**). The selected bundles were then filtered by length to avoid unrealistic streamlines and discard false positives (**6-**). To be kept in a bundle, a streamline had to be longer that half of the associated PP length, and be less than three times that length. The resulting short-range connectivity is illustrated for one example subject in (**7-**). Control PPs of each individual were analyzed with the exact same procedures.

We performed an additional analysis extracting the overall short range connectivity between the superior and middle temporal gyrus (**supplementary figure S1**). This allowed us to qualitatively assess the robustness of our method by extracting the STS connectivity without any assumption on the location of PPs and the size of seeds. Here, the entire length of the STG and MTG lines were used to define seeds, similar to the method used for the central sulcus in (Pron et al. 2018). Two seeds per hemisphere were generated by dilating the gyri lines up to 80% of the local sulcal depth in order to cover most of the gyrus walls while avoiding overlap of the two seeds. We then applied the same procedure than before to extract U-shape fibers linking the STG and MTG pair of seeds for each individual. The minimal and maximal length of streamlines were set to 20 and 80 mm respectively (Avila et al. 2019) although we tested shorter intervals into this range to observe how connectivity is affected by this parameter.

### Statistical analysis

We reported the number of PPs per individual and compared the distribution between hemispheres taking into account all PPs first, then superficial and deep PPs separately. These comparisons were performed using a non-parametric Wilcoxon signed rank test that assumes a possible dependency of PP numbers across hemispheres in a same individual.

We then tested our hypothesis of PPs being associated with a specific short-range connectivity. To this aim, we counted the number of streamlines extracted from each pair of superficial, deep or control PP seeds, for all seed sizes. We compared these three categories two-by-two in terms of their number of streamlines using a Mann-Whitney rank test. This test was performed for all size of seeds within each hemisphere, then across hemispheres.

## Results

### Plis de passage

We identified 2 to 8 PPs in the left STS (mean: 4.5) against 1 to 7 in the right STS (mean: 4.3) with bifurcations frequency of 20% in both. We found in total more PPs in the left hemisphere (total of 408) compared to the right (total of 388) but this difference was not significant across individuals (p=0.1; Wilcoxon). Between these true PPs, we were able to generate 144 control PPs in the left and 117 in the right hemisphere.

From the DPF maps we extracted one maximal DPF value per PP to plot their DPF distribution as done in (Le Guen, Leroy, et al. 2018). What stands out from this graph (**Figure 4a**) is the difference of DPF distribution between hemispheres. We can observe a greater proportion of PPs with negative or low positive DPF (i.e. superficial) in the left STS (light grey), and a higher number of PPs with high positive DPF (buried in the depth) in the right STS (dark grey). This distribution is very close to what has been reported in (Le Guen, Leroy, et al. 2018). However, in this paper the authors chose to discard PPs that have a DPF values greater than 0.42 considering them as noise. Here, we use this threshold to classify our PPs into “superficial” (DPF < 0.42) or “deep” (DPF > 0.4). Based on this classification, there were five times more deep (n=327) than superficial (n=61) PPs in the right hemisphere but only twice as much in the left (n=283 compared to n=125). These proportions differed significantly in both hemispheres (p<0.05, Wilcoxon). Between hemispheres, we also found that individual proportions of both deep PPs and superficial PPs differed significantly (p<0.05, Wilcoxon). Thus, while the absolute number of PPs does not seem to vary much from one hemisphere to another, their depth level is asymmetrically distributed with a greater proportion of superficial PPs in the left STS.

**Figure 4:**
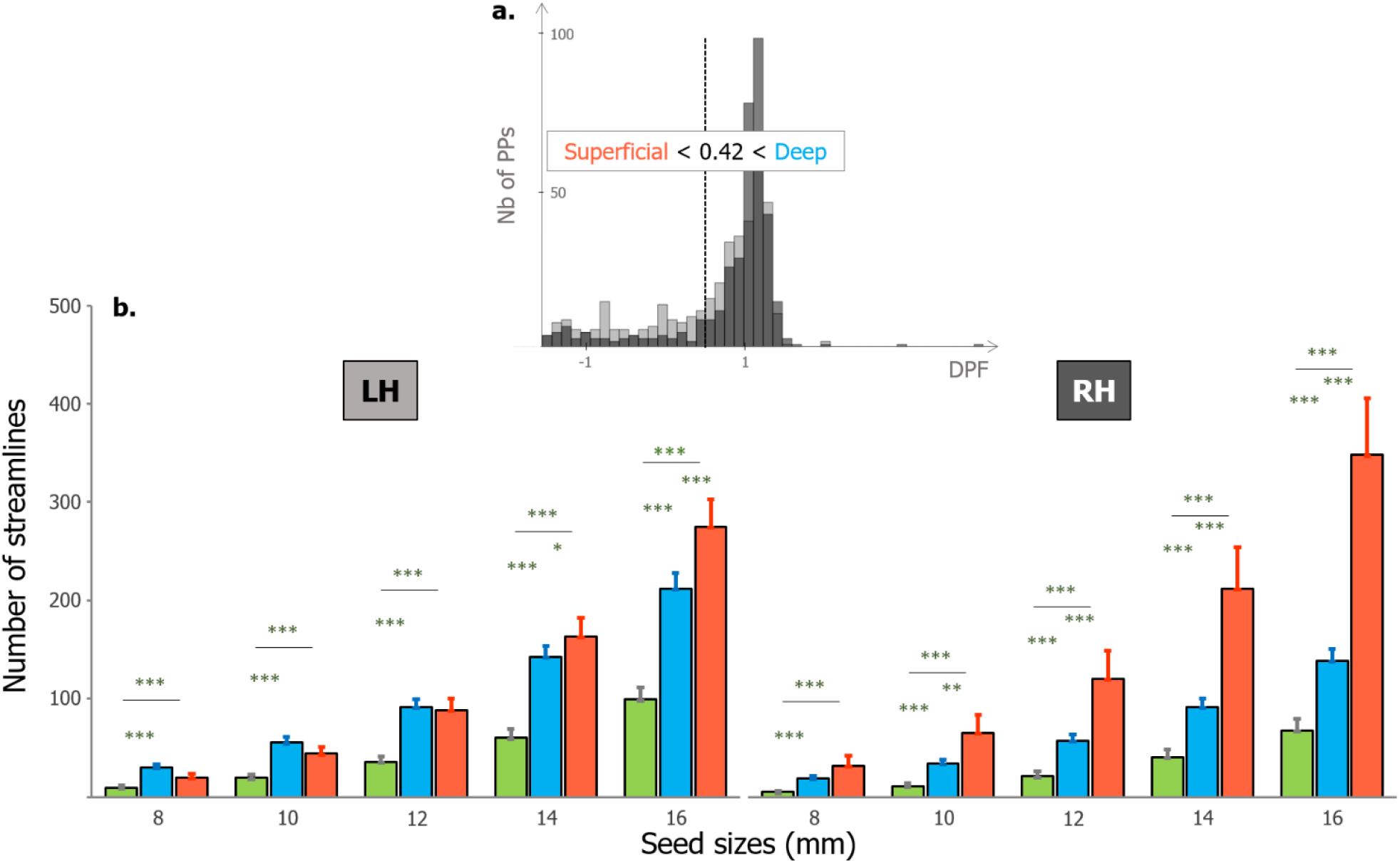
**a.** Proportion of PPs according to their DPF values for the left (light grey) and right hemisphere (dark grey). PPs were classified as “superficial” or “deep” if their DPF level was below or above 0.42 respectively (vertical dashed line), based on the threshold used in (Le Guen, Leroy, et al. 2018). **b.** Proportions of streamlines found below superficial PPs (red), deep PPs (blue) and controls (green) in the left (LH) and right (RH) hemisphere. X-axis indicates the different surfacic seed sizes used to extract streamlines (in mm). The number of streamlines increases from controls to deep and to superficial PPs for all seed sizes. Errors bars illustrate the standard error of the mean (SEM). Significant differences between categories are indicated by the stars (Mann-Whitney rank test * p<0.05 ** p<0.001 *** p<0.0001).

Some outliers emerged in the right tail of the DPF distribution as deep PPs with maximal DPF value. This is explained by our criteria used for deep PPs identification, for which an elevation of the STS fundus is not required and instead local deformations (‘pinching’) of the adjacent walls is used. Hence, outliers here can correspond to deeply buried PPs causing no STS fundus elevation (**see figure 2**, **3**).

We reported in more detail the individual patterns concerning the deep and superficial PPs (**supplementary material figure S2**). The most represented pattern in our population was “1 superficial and 4 deep PPs” in both hemispheres, with a frequency of 16% and 20% in the left and right STS respectively.

### U-shape connectivity

The main analysis of the paper examined the short-range connectivity connecting the pairs of PPs seeds. Quantitative analysis revealed an increasing number of extracted streamlines according to seed size, with significant differences between PPs and controls for all sizes (**figure 4**). The DPF-based classification into superficial and deep PPs described earlier was applied to extract the connectivity in these two categories separately. Both deep (blue) and superficial PPs (red) exhibited a higher short range connectivity (higher number of streamlines) than control regions (green) (p<0.0001, Mann-Whitney test). The quantity of streamlines obtained for superficial PPs was also significantly higher than for deep PPs, from 14 to 16 mm seeds in the left STS and from 10 to 16 mm seeds in the right STS. These results suggest that PPs are at specific locations within the STS where a majority of U-shape fibers cross the sulcus.

Because we extracted the connectivity from PP seeds, we cannot relate directly the number of PPs to that of streamlines due to circularity issues. However, we previously found that the total number of PPs in the left hemisphere did not differ significantly from that of the right hemisphere, making this question possible to assess. We found in general more streamlines in the left hemisphere compared to the right across individuals (p<0.05, Wilcoxon) for all seed sizes except 8 mm (**supplementary figure S2**). In the precedent section, we showed that the left STS contained more superficial PPs (n=125) than the right hemisphere (n=61). However, we can see in the **figure 4** that superficial PPs exhibited more streamlines in the right than in the left sulcus. Hence, the short-range connectivity appears to be more generally distributed among superficial and deep PPs in the left hemisphere, but concentrated mainly below superficial PPs in the right hemisphere.

Visual observation of the short range connectivity revealed clear short bundles that are transverse to the antero-posterior axis of the STS. Example subjects are represented on the **figure 5** for which we can compare the position of previously identified PPs and the streamlines extracted from the correspondent pairs of seeds. What emerged from the figure is that streamlines seem to co-localize with PPs positions but also that they follow the same orientation. In addition, the structural connectivity extracted from whole gyri seeds (**supplementary figure S1**) and without assumptions on PPs’ location exhibit streamlines that are not randomly organized along the STS. Instead, short bundles seem to cross the STS at the level of restricted portions distributed along the antero-posterior axis and corresponding to the identified position of PPs. On average, a greater amount of streamlines were extracted with the gyri-based method compared to those obtained from PPs seeds, across individuals and hemispheres (**figure S2**).

**Figure 5:**
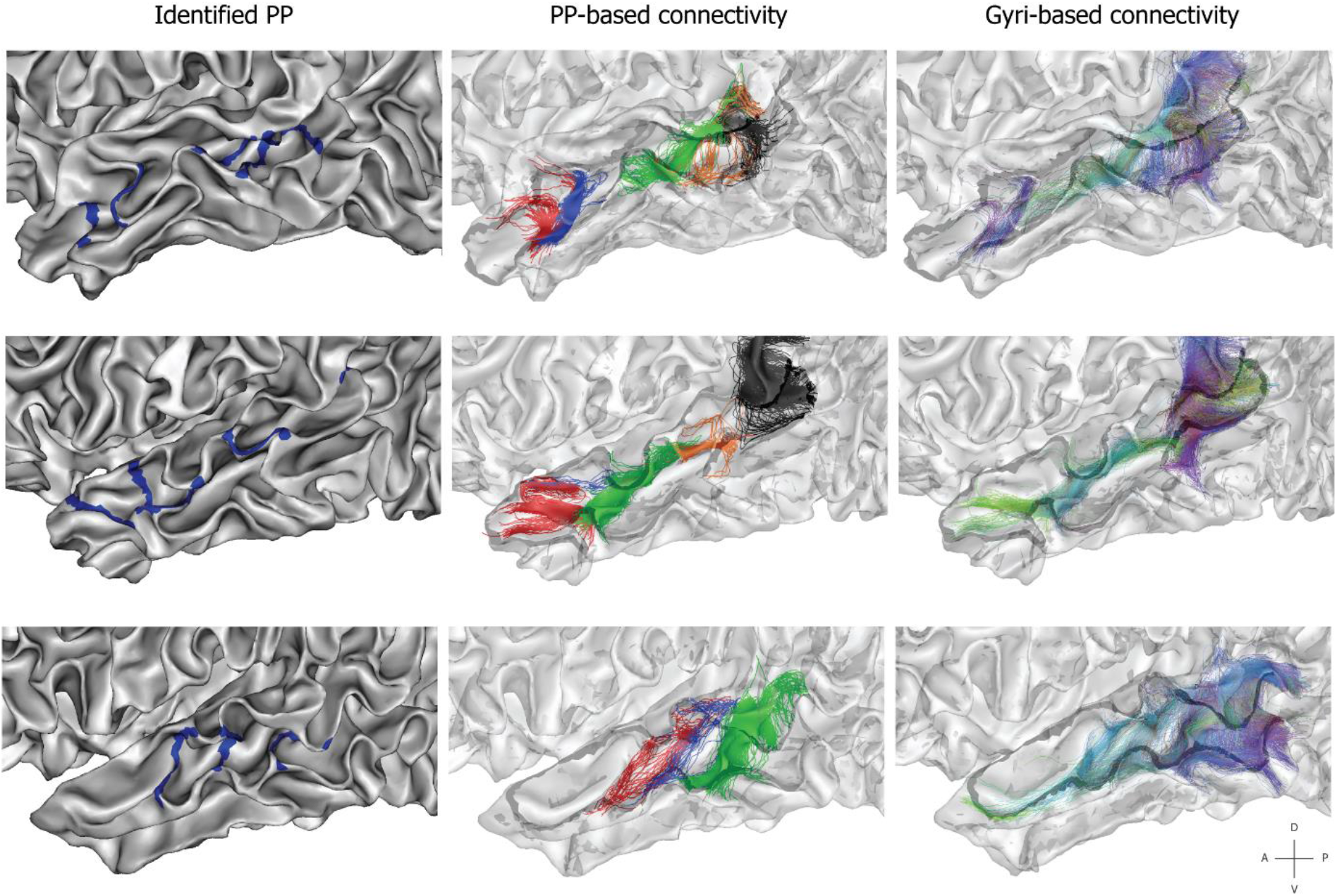
The STS short-range (U-shape) structural connectivity illustrated on three individual cortical surfaces (each row). Zoomed sagittal views of the temporal lobe are represented. **Left:** Individual PPs previously drawn on the surface (blue lines). **Middle:** Short-range connectivity extracted from PPs seeds. Each color represents the streamlines extracted from one distinct PP. The transparency of the mesh was increased for visualization. **Right:** Short-range connectivity extracted from the whole gyri seeds. A color code was attributed to each streamline depending on its orientation in relation to the STS. The black lines represent the gyri crests adjacent to the STS.

We made two additional observations from the reported connectivity patterns. First, the start and end points of the streamlines appeared generally distributed in a fan shape on the gyri walls and tighten in the middle, below the fundus of the STS like a bottleneck, as illustrated on the **supplementary figure S4** for two example PPs. Second, there was generally an increasing number of U-shape fibers from the anterior to the posterior part of the STS. This was particularly true in the case of the gyri-based connectivity (**figure 5 right**), for which streamlines are extracted further in the temporo-parietal junction. Because we stopped our identification of PPs before the intersection of the STS horizontal and posterior branches, this can also explain the greater amount of streamlines extracted from gyri seeds compared to PP seeds (**figure S3**).

## Discussion

### PPs as local and gradual cortical deformations

The results presented in this study reinforce the idea that PPs are important landmarks in the STS organization. They can be associated with specific local geometrical features, “wall pinches” from surrounding gyri (here STG and MTG), an increase in the curvature of the STS fundus and a local decrease of its depth. These criteria are in agreement with the earlier anatomical studies (Gratiolet 1854; Cunningham 1892; Ono, Kubik, et Abernathey 1990; Zlatkina, Amiez, et Petrides 2016). In particular, our observations in the STS fitted particularly well with those of Cunningham for the central sulcus: *“here is generally a shallowing of the fissure and a deep interlocking of its adjacent walls. Two of the interdigitating gyri, one projecting backwards from the anterior central convolution* [here downward from the STG]*, and the other forwards from the posterior central convolution* [here upward from the MTG] *are always larger and more pronounced than the others, and in a considerable number of cases they unite at the bottom of the sulcus in the form of a distinct deep gyrus, which constitutes a marked interruption in its floor.”* (Cunningham 1892; White et al. 1997). In addition, instead of a binary categorization of the presence or absence of PPs that eliminates the most buried ones based on their depth (Leroy et al. 2015; Le Guen, Leroy, et al. 2018), we used a method that assumed a continuum from the deepest to the most superficial PPs. Cunningham’s words are in line again with this idea: “*All gradations between a mere shallowing with an interlocking of the adjacent walls of the fissure and the presence of a distinct deep annectant gyrus are met with”*.

We believe that PPs constitute good candidates for a finer description of the STS spatial organization and morphology (Ochiai et al. 2004; Regis et al. 2005). Indeed the “sulcal roots” model provides a template organization of folding to address inter-suject correspondences (Regis et al. 2005) but still lacks flexibility, especially in the case of very complex sulci such as the STS. Based on this model, another method consists in the detection of sulcal pits (Lohmann, von Cramon, et Colchester 2008; Im et al. 2010; G. Auzias et al. 2015) which relies on reproducible and highly local depressions of the surface. Although they were shown relevant for studies on development (Meng et al. 2014; Brun et al. 2016; Im et Grant 2019) and heritability (Le Guen, Auzias, et al. 2018) we still do not know their exact relationship with morphological patterns of sulci. For this purpose, PPs may be more appropriate, as they exhibit a complex geometry dependent on the shape of surrounding gyri (**figure 2**, **3**) and closely related to the underlying white-matter (**figure 4**, **5**).

### PPs as short-range connectivity pathways

We found that PPs as we defined them morphologically constitute specific local pathways for the U-shape structural connectivity linking the two banks of the STS (**figure 4**, **5**). All PPs were considered as gradual forms of a depth continuum and identified based on specific morphological criteria (**figure 2**, **3**). In a second step, we classified them into “deep” (DPF > 0.42) and “superficial” PPs (DPF < 0.42) and both exhibited a higher connectivity than control regions were no PPs has been found (**figure 4**). This result suggests that PPs, regardless of their depth, are anatomical features of interest and that restrictive identification from the depth profile (Leroy et al. 2015; Le Guen, Leroy, et al. 2018) can miss crucial information. Comparison of the results obtained for PPs- and gyri-seeds reinforced our hypothesis of a dense U-shape connectivity below PPs locations. Instead of a random distribution of fibers between the STG and the MTG, we clearly observed several bundles along the antero-posterior axis of the STS whose location appears to correspond to that of PPs (**figure 5**).

The results we obtained are in agreement with the U-shape connectivity found along the central sulcus (Catani et al. 2012; Magro et al. 2012; Pron et al. 2018), but differ from recent investigations of the short-range connectivity in the STS (M. Guevara et al. 2017; Abouzahr et al. 2019). Indeed, these authors reported only one bilateral bundle in the posterior STS that was orthogonal to the sulcus and connecting the STG and MTG. This discrepancy could be attributed to the fact that these studies aimed to extract only the most reproducible bundles across individuals whereas we have considered all the streamlines under PPs here. Nevertheless, this could suggest that this posterior bundle is more genetically constrained, or at least more systematically present, compared to the others. Recently, a higher heritability was also reported in this region regarding the STS depth and the presence of PPs in the left hemisphere (Le Guen et al. 2019). This posterior PP would also appear first during the development in-utero, followed by the others until the most anterior one between the 5^th^ and 7^th^ month (Ochiai et al. 2004).

Further work is required to confirm the link between short-range connectivity and PPs through their respective reproducibility in the population. In particular, it is certainly closely related to the location of the sulcal pits and must therefore be evaluated in relation to the heritability of these landmarks (Le Guen, Auzias, et al. 2018; Le Guen et al. 2019). Another approach would be to compare the occurrence of PPs and short range bundles across species. In Zhang et al. (2014), they did not found U-shape connections along the chimpanzee STS but some of them in the posterior STG of macaque monkeys, similar to what has been reported in a human atlas (M. Guevara et al. 2017). Although very few studies investigated U-fibers in primates, the presence of PPs in these species has been documented since the seminal studies (Gratiolet 1854; Cunningham 1890c). Leroy et al. (2015) noted the presence of PPs in the STS of chimpanzees but they were less numerous and more symmetrically distributed compared to humans (again this result should be modulated by the PPs detection method). Inter-species comparison raises interesting questions about the functional role of PPs but also about their relationship to brain size and the level of gyrification. In light of the present results, less gyrified species may constitute ideal models for testing the association between PPs, gyrification and connectivity.

### Anatomical implications

The relation suggested here between local cortical morphology and the underlying white matter can be related to recent advances in cortical development research (Llinares-Benadero et Borrell 2019). Folding would be initially constrained by genetic factors that regulate cellular assemblies to differentiate sulci versus gyri as well as regional patterns. This process could also determine some crucial tissue properties of the future cortex such as the stiffness and thickness of its layers. These tissue properties have been shown to interfere with the tangential growth of the cortex by determining the final aspect of folding such as the wavelength of the folds at the regional scale (Bayly et al. 2013; Budday, Steinmann, et Kuhl 2014). The sulcal roots of the STS were shown to appear separately along a caudal to rostral gradient (Ochiai et al. 2004), each of them characterized by one pit separated by two transverse plis de passage (Regis et al. 2005). We can imagine that tangential expansion of the surrounding STG and MTG portions follow the same timeline, gradually revealing the STS valley. Orthogonal to the sulcus, this expansion would lead to a digging of the STS excepted at the location of PPs that are also expanding in the opposite direction. Parallel to the sulcus, this expansion would then induce a deformation of the PPs depending on the surrounding growing pattern. This hypothetical order of events comes from the apparent deformation of superficial PPs, often forming “S” or “C” shapes with a slight shift of the extremities on the gyri walls.

The first question that appears here is how deep and superficial PPs are formed. Especially, what are the factors that could lead to a clear interruption of the sulcus or to “wall pinching” configurations with no real depth variation? We propose here a three-step hypothesis following the developmental time scale: first, tangential growth during cortical development in utero would favor the transition from initial superficial PPs to more buried forms. This corroborates the findings of an increasing sulcal depth (Glasel et al. 2011) and fusion phenomenon of sub-parts of the folds during this period (Cunningham 1892). The early maturation of the right STS (Glasel et al. 2011; Rajagopalan et al. 2011) would result in prolonged erosion of PPs giving rise to a deeper, less interrupted sulcus compare to its left counterpart (Ochiai et al. 2004; Glasel et al. 2011; Leroy et al. 2015). Then, a second step would involve the gradual introduction of U-shape fibers joining the STS walls, passing in larger proportions through the most direct trajectories provided by superficial PPs, as previously suggested (J.-F. Mangin et al. 2019). We have shown in this respect fewer streamlines under deep PPs and a generally lower proportion in the right (and deeper) STS (**figure 4**). It is important to note here that the aforementioned short-range connections seem to appear late in development, around birth and up to 6 months after (Kostović, Sedmak, et Judaš 2019). Hence, the third step would occur postnatally and involve in particular the later expansion of the cortex. The U-shape connectivity would serve here as a holding scaffold maintaining the pre-established PPs. More precisely, the weak (or strong) connectivity under the preexisting deep (or superficial) PPs making them more (or less) vulnerable to the surrounding expansion and hence lead to a “wall pinching” remnant (or superficial bridge). This secondary interaction between connectivity and PPs may be more exposed to environmental factors and potentially explain the large variability observed across individuals. Indeed in some subjects we identified more PPs (up to 8 in the left STS) than assumed by the sulcal roots model (Regis et al. 2005; Ochiai et al. 2004). The functional properties inherent to each PP regions could be an important factor as well, especially in the determination of individual patterns (J.-F. Mangin et al. 2019).

Together, these considerations raise the need for novel longitudinal studies using in-utero imaging that would specifically follow the evolution of each PP, as already done for the sulcal pits (Meng et al. 2014) or folding patterns (Duan et al. 2018).

### Functional implications

There are two main functional implications of PPs, which are related to their asymmetry and spatial organization. On one hand, the asymmetric distribution of PPs could be related to functional asymmetry of the temporal region. We found that the left STS exhibited a stronger U-shape connectivity than the right while their respective number of PPs did not differ significantly. There are at least two possible explanations for this result. The most obvious concerns the greater proportion of superficial PPs in the left STS, which confirm earlier observations (Ochiai et al. 2004). As we found generally more streamlines under superficial than deep PPs, the connectivity in left STS would be consequently higher. However, although the difference in number of streamlines is large between deep and superficial PPs in the right STS, it is less so in the left STS (**figure 4**). Another possible explanation could be linked to other asymmetric properties of the STS. It is well know that language related functions are strongly lateralized in the brain, however, this mostly involves the fronto-temporal networks (Hickok 2012). Nevertheless, some specializations exist between the two temporal lobes such as temporal processing in the left and spectral processing in the right auditory cortex (Zatorre et Belin 2001).

On the other hand, several studies attested to the relevance of PPs for functional localization. The position of sulcal interruptions was used to explain those of functional activity in various regions such as the intra-parietal (Zlatkina et Petrides 2014), cingulate (Paus et al. 1996; Amiez et al. 2013) and post-central (Zlatkina et Petrides 2010; Zlatkina, Amiez, et Petrides 2016) sulci. In the fusiform gyrus, the presence of one PP in the visual word form area was recently associated to better reading skills, and length of the interruption correlated positively with these ability (Cachia et al. 2018). In the central sulcus, the PPFM co-localize with the hand motor area (**figure 6d.**) (Boling et al. 1999; Cykowski et al. 2008; J.-F. Mangin et al. 2019) and with U-shaped fibers bundle (Magro et al. 2012; Pron et al. 2018). This correspondence is sufficiently robust that a variation in the position of the hand-knob is followed by that of the motor activation along the central sulcus (Sun et al. 2015). Such relation was never investigated for the STS PPs, mainly because of the complex functional organization of the STS, also because interruptions vary to a greater extent across individuals. However, functional studies can help to identify candidate functional areas that could be associated with the STS PPs.

**Figure 6:**
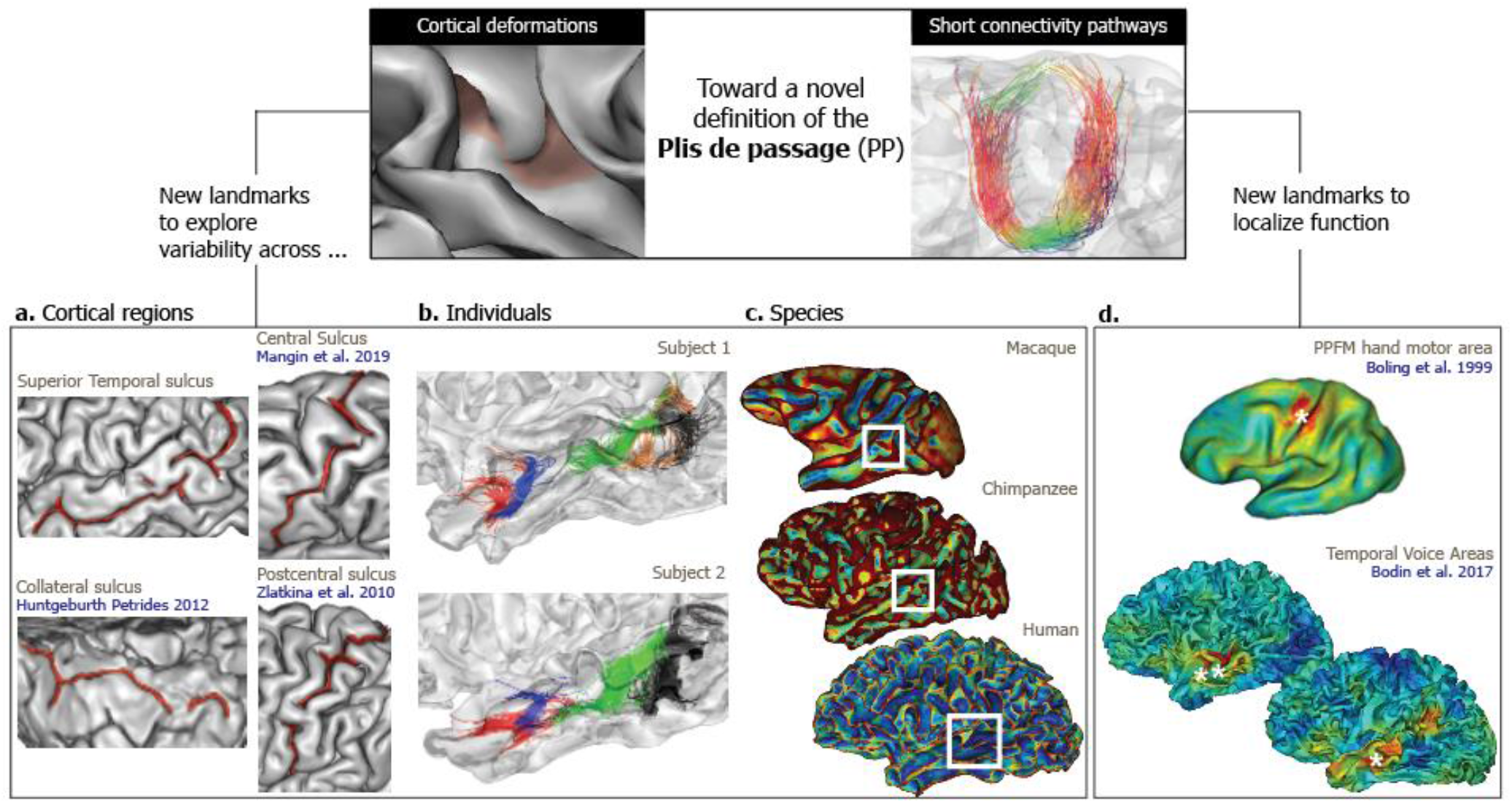
Potential use of plis de passage (PP). **a.** Example sulci (red) labelled and extracted automatically in BrainVISA with a clear PP visible at the pial surface. **b.** Two example subjects of the present study showing a similar organization of their PP-associated connectivity. **c.** Curvature maps of three primate species projected on their respective white mesh (generated in BrainVISA). The white rectangle indicates a possible PP characterized by a local increase of the curvature and wall pinching. **d.** Individual functional maps of the hand motor area in the central sulcus (up, see panel a.) or voice areas in the STS (below) suggest a tight relationship with the location of PPs in these sulci (white stars). Red color indicates a stronger functional activity.

As previously noticed, we observed a dense connectivity in the pSTS that is often constituted of several bundles connecting close territories. This could be related to the complex functional organization in the region encompassing the pSTS and the temporo-parietal junction (TPJ) (Patel, Sestieri, et Corbetta 2019). On the right hemisphere, this region was shown to hold high-level social functions such as the representation of identity by integrating both facial and vocal information (Watson et al. 2014; Hasan et al. 2016; Davies-Thompson et al. 2018; Tsantani et al. 2019). In the left hemisphere, this region was mainly associated to the dorsal pathway of language implicated in articulatory and production mechanisms (Hickok et Poeppel 2004). In the middle temporal region, Upadhyay et al. (2008) evidenced two streams going from the primary auditory cortex toward the anterior and the posterior STG using effective connectivity, and these areas where also found connected by structural fiber pathways. Here, we observed several cases of bifurcations (see methods) crossing the middle STS that could correspond to this streams, as on line 2 of the **figure 5**.

In a previous study (Bodin et al. 2017), we found a correspondence between the local depth maxima in the STAP region (Leroy et al. 2015) and the voice areas’ (Belin et al. 2000) maximal activity. At the individual level, this relation was less systematic notably because of the variable position of voice-related activity and depth pits along the STS. Interestingly, in many cases we observed a maximal activity not in the depth pit but on one of the PPs adjacent to it (**figure 6 d.**), which were already identified using the depth profile method on both sides of the STAP (Leroy et al. 2015; Le Guen, Auzias, et al. 2018; Le Guen, Leroy, et al. 2018). The fact that voice-related activity can be clustered into three areas from the anterior STG to posterior STS (Pernet et al. 2014) and that these areas are functionally inter-connected with each other’s (Aglieri et al. 2018) opens numerous questions on their link with PPs and therefore with the U-shape structural connectivity.

### Limitations of the study

This study was based on the manual identification of PPs based on the cumulated expertise of C.B. and from a previous investigation on the STS (Bodin et al. 2017). Thus, we assume that this method may have included a proportion of noise in the PP identification procedure. In particular, the fact that deep PPs are more difficult to identify and were more numerous in the right STS could have participated to the difference in the number of streamlines found between superficial and deep PP in the right hemisphere (**figure 4b**).

New detection tools are needed to test the strength of the association between local morphology and connectivity while avoiding time-consuming and operator dependent manual steps. In addition, streamlines extraction could be improved to prevent the overlap of seed regions. Indeed, in the case of two close neighboring seeds A and B (because of close PPs), it happened that streamlines extracted from A actually passed through the PPs corresponding to B. As mentioned above, although the streamlines were tightened in the STS, their extremities were more spatially dispersed on the gyri walls (**figure S4**). Thus, future studies should also take into account the trajectory of streamlines according to that of PPs crests.

One general limitation concerns the plausibility of the results obtained with diffusion tractography. A recent report showed that the tractograms available to date contain more invalid than valid bundles in comparison to ground truth (post-mortem based) studies (Maier-Hein et al. 2017). In particular, regions that are exhibiting multiple bundles (called “bottlenecks”) such as the temporal lobe can lead to spurious tractographic reconstructions. Nevertheless, in this report they did not describe U-shape short-range connectivity. Here, we used a similar method to Pron et al. (2018) in which U-shape fibers were found under the PPFM location in the central sulcus, and already describes in another study relying on post-mortem blunt dissections (Catani et al. 2012). The method is based on state-of-the art tractography methods followed by a filtering step (COMMIT (Daducci et al. 2015)) that reduced the number of streamlines by 80%, securing those that explains best the signal in dMRI images. We also filtered streamlines based on their length to add some anatomical plausibility. These arguments advocate in favor of a probable U-shaped connectivity under the STS PPs, although further work is needed to assess its robustness.

## Conclusion and potential use of PPs

This study showed that morphological criteria identifiable from individual cortical surfaces can be used to detect and characterize the plis de passage. This lead us to suggest that the previously described asymmetry in the number of PPs between the left and right STS might in fact be an asymmetry of their depth but that their number is similar between both hemispheres. We demonstrated, for the first time in the superior temporal region, the nature of PPs as key places where short white-matter fibers converge to cross the STS. This heterogeneous fibers distribution appears to be modulated by the depth level of the PPs, with higher connectivity observed below the shallowest ones. Importantly, we do not claim to provide a definitive definition of PPs, rather, we suggest reconsidering these structures as continuous variations of morphological features and the U-shaped local connectivity as a new marker for the presence of PPs in the STS (**top of figure 6**). These landmarks open up new avenues of research for a more detailed description of cortical anatomy across different cerebral regions and a new index to investigate individual variability. Further research might also explore whether PPs are present in other species, indeed, the cortical surface in monkeys and chimpanzees seems to present morphological deformations similar to those of deep PPs in humans (**figure 6 c.**). New investigations are needed to characterize them in light of the current results on U-shape connectivity. Finally, PPs could provide landmarks to localize the functional activity as done in the central sulcus or to reduce the individual variability typical of high-level functions (**figure 6 d.**).

## Acknowledgements

This work was supported by grants ANR-16-CONV-0002 (ILCB), ANR-11-LABX-0036 (BLRI) and the Excellence Initiative of Aix-Marseille University (A*MIDEX).

