## supplementary material for "Plis de passage in the Superior Temporal Sulcus: Morphology and local connectivity"

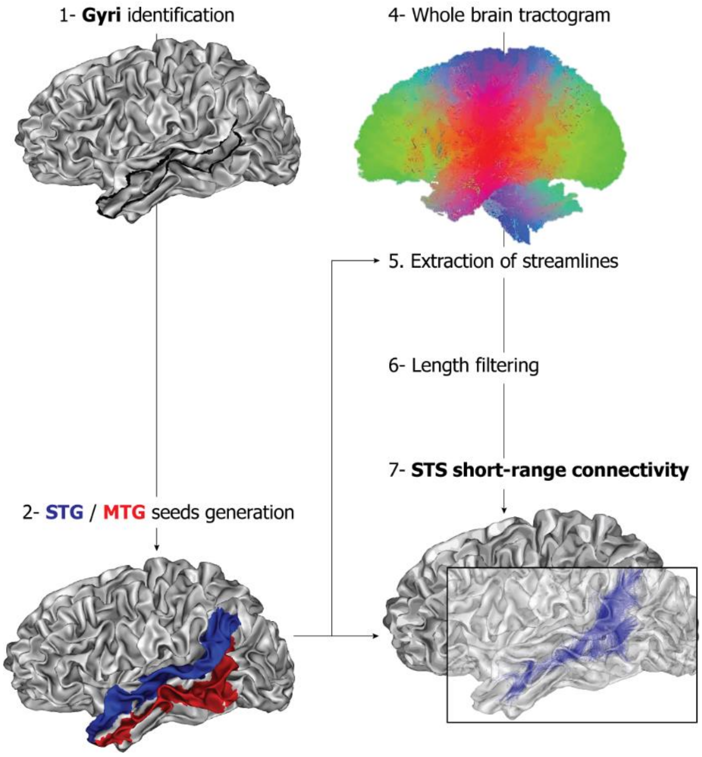


**Figure S1:** Additional analysis where gyri lines (1-) served to build bigger seeds (2-) corresponding to the STG (blue) and the MTG (red). Streamlines connecting these two seeds were then extracted without any constraint of PPs locations (5) and represent the overall short range connectivity of the STS.

| LEFT | | Number of superficial PPs | | | | |
|---|---|---|---|---|---|---|
| | | 0 | 1 | 2 | 3 | 4 |
| Number of deep PPs | 0 | 0 | 0 | 1 | 0 | 1 |
| | 1 | 0 | 0 | 7 | 2 | 0 |
| | 2 | 0 | 8 | 12 | 3 | 0 |
| | 3 | 2 | 11 | 9 | 2 | 0 |
| | 4 | 7 | 16 | 3 | 0 | 0 |
| | 5 | 4 | 4 | 1 | 0 | 0 |
| | 6 | 1 | 3 | 1 | 0 | 0 |

| RIGHT | | Number of superficial PPs | | | | |
|---|---|---|---|---|---|---|
| | | 0 | 1 | 2 | 3 | 4 |
| Number of deep PPs | 0 | 0 | 0 | 0 | 0 | 0 |
| | 1 | 1 | 3 | 1 | 0 | 0 |
| | 2 | 1 | 8 | 2 | 0 | 0 |
| | 3 | 4 | 18 | 2 | 0 | 0 |
| | 4 | 14 | 20 | 1 | 0 | 0 |
| | 5 | 16 | 4 | 0 | 0 | 0 |
| | 6 | 2 | 1 | 0 | 0 | 0 |

**Figure S2:** Frequency of subjects (%) exhibiting similar PPs patterns for the left (upper table) and right (lower table) hemisphere. For each individual we reported its number of deep and superficial PPs in the STS. Colors code the increasing frequency of subject having a similar pattern from lightest to darkest. The maximal number of PPs observed was 8 and 7 in the left and right STS respectively.

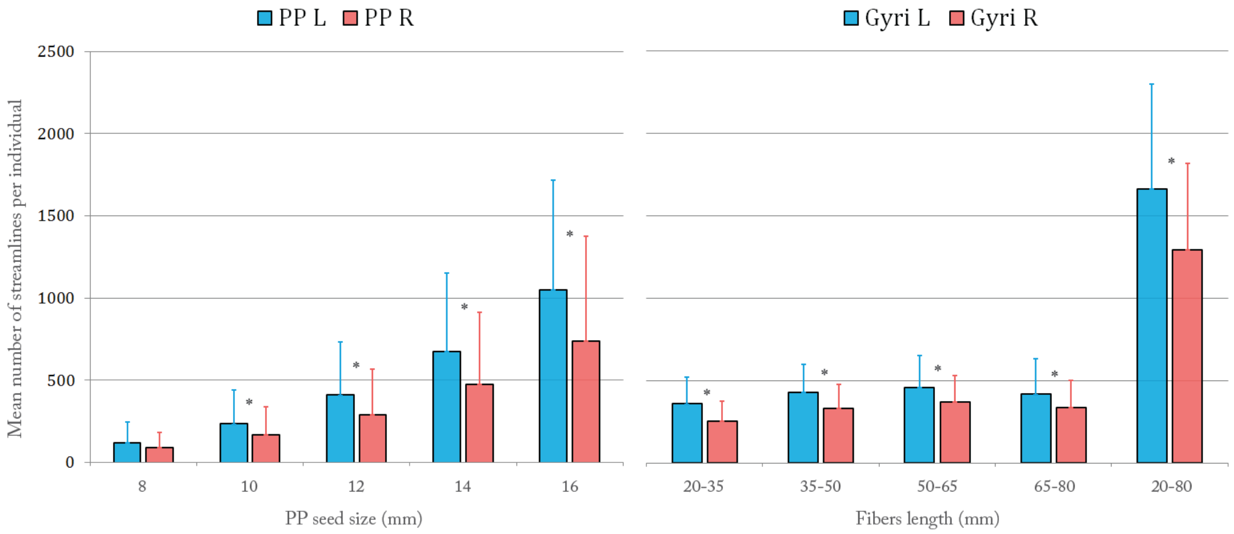


**Figure S3**: Average number of streamlines found in individuals from all their PPs seeds (left panel) and the gyri seeds (right panel) in the left (blue) and right hemisphere (red). The mean number of streamlines is plotted according to PPs seed sizes and the range of streamlines length (in mm) for the gyri seeds. All the 15 mm long ranges showed more streamlines on the left than on the right but differed very slightly between them. By filtering fiber length between 20 and 80mm, however, we extracted on average 1663 streamlines on the left and 1294 streamlines on the right from one individual STS. This is higher than the average number of streamlines extracted from individual PPs using the biggest seed size (16 mm). Stars indicate significant difference between left and right proportions (Wilcoxon rank test).

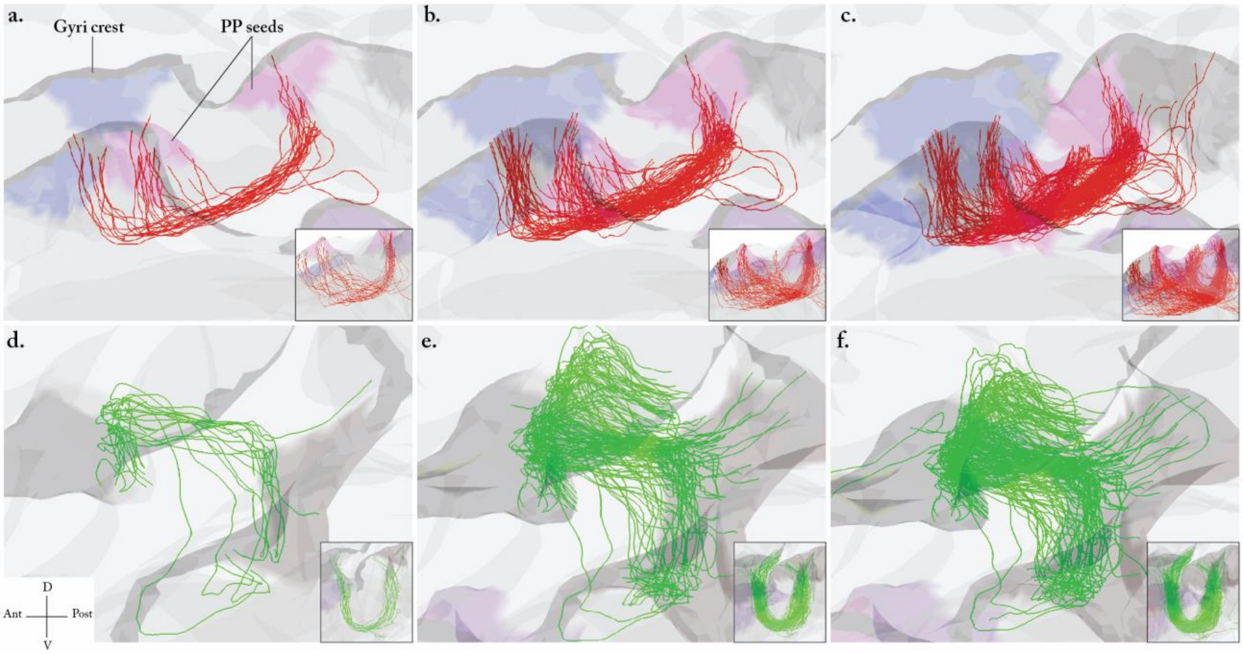


**Figure S4**: Illustration of streamlines converging through PPs pathways. Streamlines extracted from two example PPs (a-c and d-f) represented from the sagittal view of the individual mesh (main panels) for different seed sizes: 8 mm (a,d), 12 mm (b,e) and 16 mm (c,f). A lateral view is also plotted to reveal the U-shape connectivity (attached small panel).
